# The Lair: A resource for exploratory analysis of published RNA-Seq data

**DOI:** 10.1101/056200

**Authors:** Harold Pimentel, Pascal Sturmfels, Nicolas Bray, Páll Melsted, Lior Pachter

## Abstract

Increased emphasis on reproducibility of published research in the last few years has led to the large-scale archiving of sequencing data. While this data can, in theory, be used to reproduce results in papers, it is typically not easily usable in practice. We introduce a series of tools for processing and analyzing RNA-Seq data in the Short Read Archive, that together have allowed us to build an easily extendable resource for analysis of data underlying published papers. Our system makes the exploration of data easily accessible and usable without technical expertise. Our database and associated tools can be accessed at The Lair: http://pachterlab.github.io/lair

## Background

The Short Read Archive (SRA) is a public repository for sequencing data that has become an important archival resource for reads associated with published papers. The accumulation of large amounts of data in the SRA allows for meta-analyses that are possible only thanks to the centralized, open sharing of data by multiple investigators [1,2]. Reads in the SRA also allow, in principle, for the reproduction of results in publications [3]. However the bioinformatics difficulties associated with processing and analyzing sequencing data [4] have limited the utility of the SRA and have made it prohibitive for most investigators to perform exploratory data analysis (EDA) on the data in the archive. The use of the SRA for EDA is especially difficult for RNA-Seq data. This is because even the most basic processing of RNA-Seq reads requires numerous decisions about appropriate software to use, complex choices about annotations, understanding of experimental design, and frequently, significant computational resources. As a result, workflows designed for operating on multiple datasets in the short read archive have mainly been restricted to the tasks of aligning and quantifying reads [5]. Similarly, the Gene Expression Omnibus (GEO) requires an expression matrix to be uploaded with every sample, however these are static and frequently out of date, and fail to provide users with the complete analyses that they typically seek to explore.

We have recently developed a pair of tools called kallisto [6] and sleuth for RNA-Seq analysis that address a number of the challenges associated with processing RNA-Seq data. The kallisto program circumvents the need for large alignment files, a convenience that reduces storage needs and increases speed, thus enabling the processing of large numbers of samples on modest computational resources. The program sleuth utilizes bootstraps output by kallisto for differential analysis, and provides a complete interactive and web-compatible solution for exploring and analyzing RNA-Seq data. This has allowed us to semi-automate the process of associating interactive Shiny-based [7] websites for EDA of RNA-Seq data.

The processing underlying the resource we have developed can easily be rerun, providing a scalable and updatable push-button system for the analysis of large numbers of datasets. Thus, we have been able to create a semi-automated system for analysis of archived RNA-Seq data that is much more informative than mere alignment and quantification and that opens up the analyses of published data.

## Construction and content

The infrastructure for The Lair is based on Snakemake [8], a python-based workflow system. The system is organized as shown in Figure 1 and consists of three main parts: (1) an initial processing of data to produce sleuth objects that are deployed in a Shiny database; (2) a Shiny server and database; and (3) a website that can be constructed automatically and that links to the Shiny server.

**Fig. 1.**
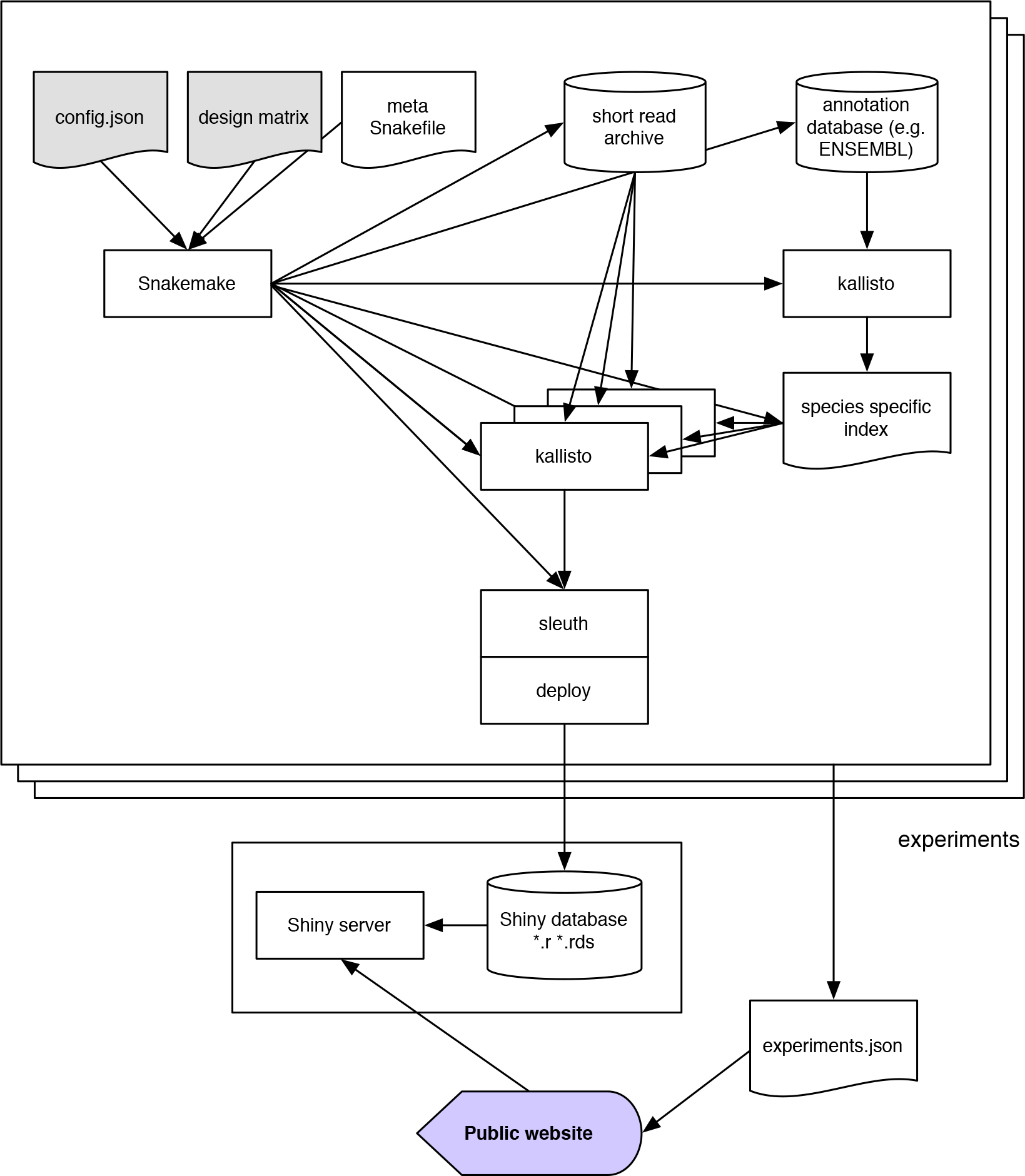
Workflow of The Lair system for distributing analysis of short read archive data. The inputs to the system are sets of two files: config.json file that specifies parameters to be used during the processing of each experiment and a design matrix for each experiment that specifies its structure. A master Snakemake workflow organizes a series of computations starting with downloading of data to the short read archive and ending with deployment of a sleuth analyses to a Shiny server. Finally, a website generated from information in the config.json files links to objects in the Shiny server thus providing access to the processed experiments.

The specification for datasets is stored on GitHub at https://github.com/pachterlab/bears_analyses. Each dataset requires a config.json file that provides information about the dataset that will be used during its processing and deployment. For example, the config.json file for a recent paper on a HOXA1 knockdown transcriptome survey in human [9] is:

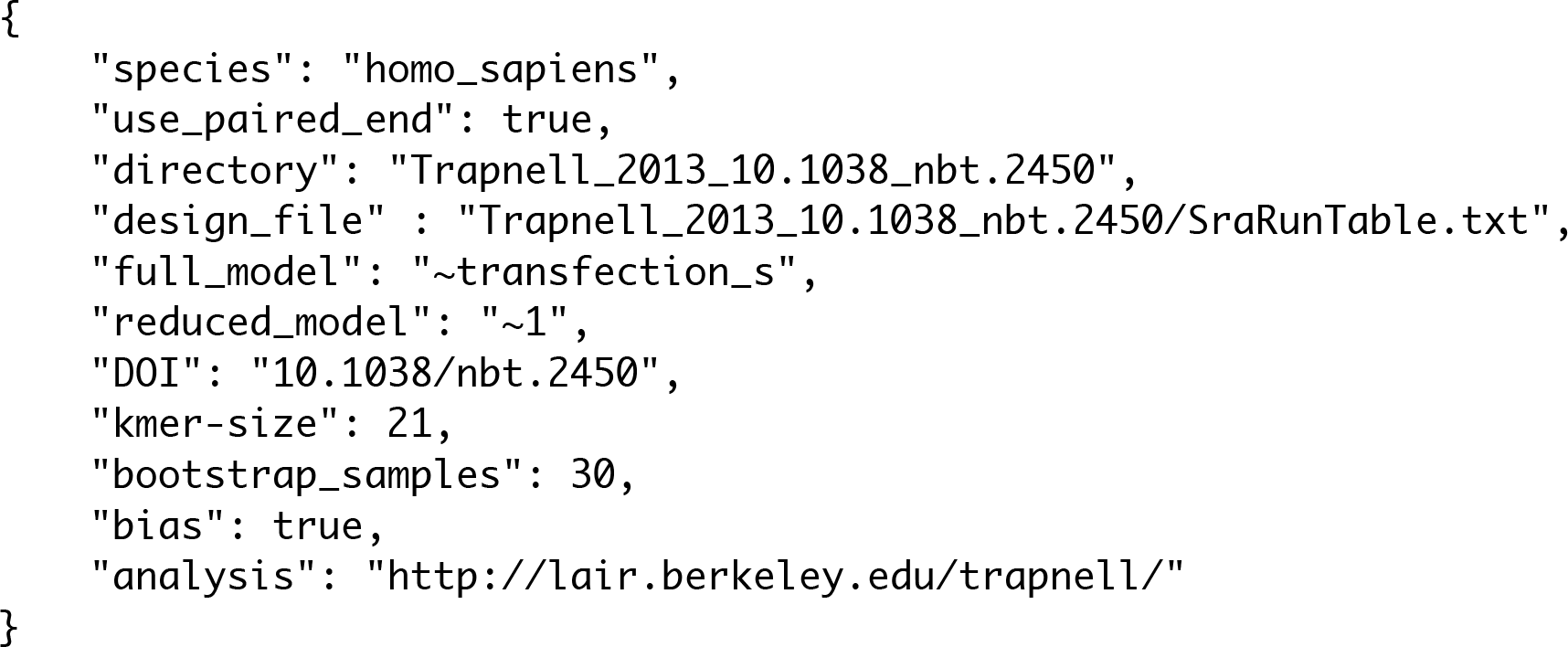

The config.json file specifies the species name for the RNA-Seq analysis, thus allowing for automatic downloading and indexing of the appropriate Ensembl [10] transcriptome for the analysis, the type of reads (single or paired), the design matrix and type of testing to perform, as well as the DOI of the paper and parameters for the kallisto processing. Along with the config.json file, a design matrix must be included which specifies the structure underlying the samples. The design matrix can be downloaded from the SRA but sometimes requires manual curation due to inconsistencies in SRA formatting. The entire workflow begins with only these two files, from which all the relevant information is extracted for processing.

The organization of The Lair allows for updating of all the analyses at the push of a button. This is useful in the case of updates to the component programs and the emphasis on speed of the constituent software allows for frequent updating. In fact, the current main bottleneck with The Bair’s Lair is downloading of the SRA data after which the entire workflow processes individual samples in minutes [6].

The website is a static page built using Jekyll [11] and Bootstrap v3.3.6 [12]. The Analyses section of the website contains a dynamically generated table of paper with corresponding live analyses. The table is powered by the JQuery plug-in DataTables [13] and features filtering and sorting by Authors, Title, Journal and Date. The title of each paper links to the original paper published in the stated journal. If the paper was published in print, the date given is the paper's print date; otherwise, the date is the paper's online publication date. The analysis button links to an in-browser analysis of the experiment, made possible by sleuth's efficient use of statistical bootstrapping and RStudio's Shiny plug-in; the analysis for each paper is generated automatically by the above build system.

The table is populated automatically by data from the bears analyses (https://github.com/pachterlab/bears_analyses) Github repository. This means that anyone can submit additional datasets for processing via Git pull requests which once accepted will become part of the website.

## Utility and discussion

To demonstrate the utility of The Bair’s Lair we examined the results from our analysis of the Trapnell *et al.* [9] data. In that paper, an RNA-Seq differential analysis was performed on lung fibroblasts responding to the knockout of the developmental transcription factor HOXA1. First, it is easy to confirm that The Lair analysis replicates the main results of the paper. Figure 2 shows a principal component analysis of the data, confirming high quality data with substantial separation between the two conditions.

**Fig. 2.**
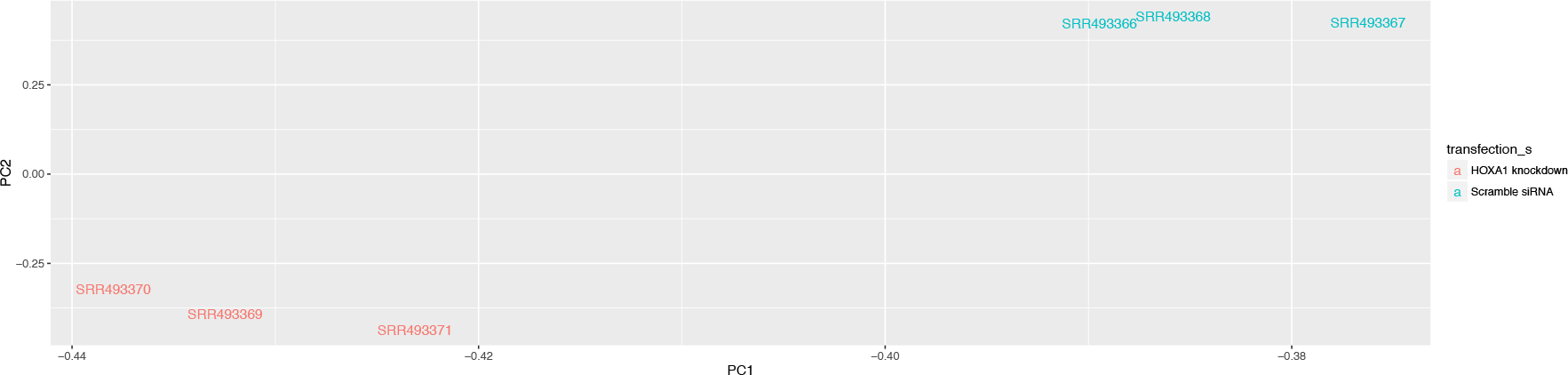
Principal Components Analysis of the Trapnell *et al.* HOXA1 knockdown RNA-Seq data. The Lair allows for plotting projections with respect to any pair of principal components, and also identifies the transcripts constituting the loadings of each dimension.

Specific results about individual genes that are discussed in the Trapnell *et al.* paper are easily confirmed. For example, Figure 5 shows transcript abundance changes in response to the knockdown, providing examples of some key genes of interest. The T-box DNA binding domain TBX3 displays an increase in exactly one out of three isoforms (Figure 5d). The differential isoform, ENST00000349155, is displayed via the “transcript view” feature in the Shiny app as shown in Figure 3. The associated q-value in the sleuth test is q=3.15e-05. The two remaining isoforms of the gene are not significantly differential, in concordance with the Trapnell *et al.* paper (for ENST00000257566, q = 0.148 and in the case of ENST00000613550, the isoform did not pass the requisite filters to be tested).

**Fig. 3.**
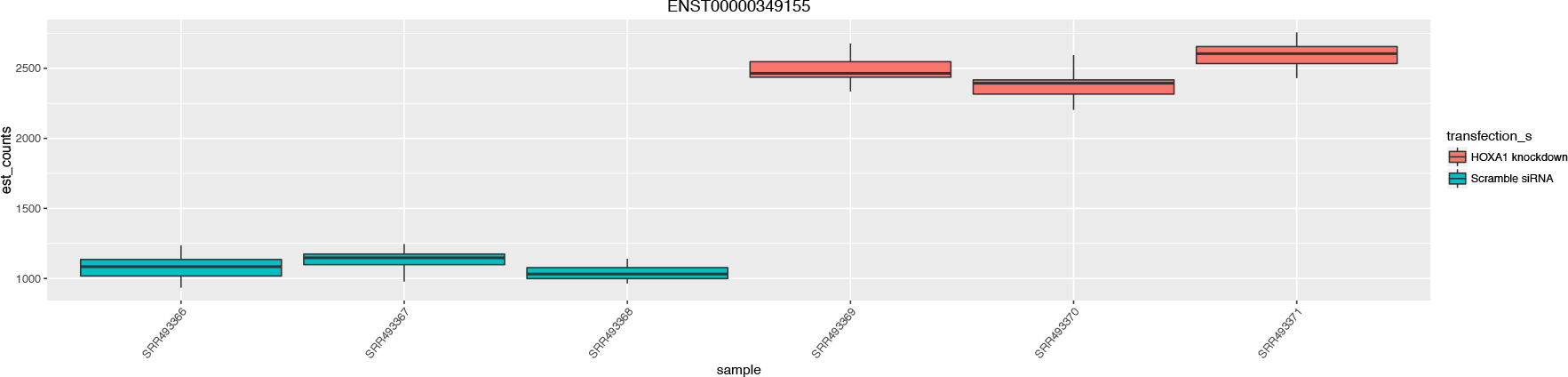
Transcript abundances for the differential isoform of the TBX3 gene in the Trapnell *et al.* data. The error bars on each quantification are produced via the bootstrap feature of kallisto, which establishes the inferential variance associated with quantification. The Lair provides an interactive template for viewing such plots for any transcript.

While the reproducibility of results is reassuring, the innovation in The Lair resource is the ability to go beyond the limited view of the data provided by the authors. In the Trapnell *et al.* example there are thousands of significantly differential transcripts, and The Lair allows for viewing the raw quantifications underlying each of them. Another advantage is the ability to examine results of different papers analyzed with the same framework. For example, the Ng *et al.* [14] paper is immediately established to have much higher variance in the estimates due to having fewer replicates in each condition (two instead of three), and the common framework underlying its analysis provides a quantification of that assessment.

## Conclusion

RNA-Seq technology provides rich and complex data for analysis in projects where expression dynamics are of interest. While investigators are eager to squeeze every bit of information out of their data, there are a number of reasons why they are unlikely to be able to do so at the time of publication of their work: analysis methods and tools improve over time and data may be revealed to be useful for applications not considered at the time of acquisition.

The Lair resource we have developed opens up large volumes of RNA-Seq data for both general and targeted exploration. The modular and automated construction of our system will allow us to upgrade it over time, adding functionality and analyses as we improve and expand the kallisto and sleuth methods and programs. An added benefit of our holistic analysis of SRA data is that our use of the same tools to process diverse datasets also allows for comparison of results across studies. Future plans for The Lair include facilitating such cross-study comparisons.

While we have focused The Lair on RNA-Seq data, the ideas and tools developed in this work should be adaptable to other data types. Hopefully such work will establish tools that create symbiotic rather than parasitic relationships between data generators and data analyzers.

## Methods

### Data handling

There are two data bottlenecks in processing of short read archive RNA-Seq data: first, the downloading and storing of large read files, an issue we do not address in this paper but that can be ameliorated with compression schemes. Second, sleuth utilizes statistical bootstraps generated by kallisto as part of its differential analysis and these can be time consuming to transmit via the web. To make sleuth usable via The Lair, the code was refactored to pre-compute variance and quantile data that are sufficient and necessary statistics to generate the plots in the online visualization. The individual bootstraps estimates can then be discarded. This reduced the size of the analysis objects by orders of magnitude and allowed sleuth analyses to be shared online and loaded in standard web browsers.

### The Snakemake workflow

The Snakefile used to generate the analysis requires two input parameters: a json configuration file and a design matrix file. The configuration file has the following required parameters:

- species: species used in the experiment.
- used_paired_end: true if the experiment was paired-end, false otherwise.
- directory: the directory to put the results into.
- design_file: the name of the required design matrix file.
- full_model: formula which describes the full or alternative model used in differential analysis.
- reduced_model: formula which describes the reduced or null model
- DOI: the digital object identifier of the publication.

The configuration file also accepts the following optional parameters:

- kmer-size: the k-mer length used to build the kallisto index (defaults to 31)
- bias: perform sequence-specific bias correction during quantification (defaults to True)

The design file must be a.tsv file, one column of which is titled 'run' or 'Run_s' and contains the SRR accessions for each run in the experiment. Furthermore, the full_model and reduced_model in the config file must match column names in the design file for the differential analysis to work correctly.

The build system checks whether the FASTA reference file for the input species is already locally accessible. If not, it downloads it to the FASTA file from Ensembl. It then uses this FASTA file to build a kallisto index given the input k-mer size if the index does not already exist, and makes this index locally accessible.

Then the build system uses the 'run' or 'Run_s' column of the design matrix to download the raw SRA data and quantifies the raw data using kallisto and the index for the specified species. Once kallisto finishes quantification, sleuth is run with the likelihood ratio test using the specified full_model and reduced_model parameters. The build system then deploys the resulting sleuth analysis onto the server, where it is available to explore online.

## Availability of data and materials

The main user website is at http://pachterlab.github.io/lair. The workflow software is at https://github.com/pachterlab/bears_analyses and the website code is at https://github.com/pachterlab/lair.

## Acknowledgements

We thank the members of the Pachter Lab for contributing design and feature ideas and for suggesting the initial datasets with which to prototype The Lair. HP and LP were partially supported by NIH grants R01 HG006129, R01 DK094699 and R01 HG008164.

